# Imaging the immune synapse: three-dimensional analysis of the immune synapse

**DOI:** 10.1101/2023.02.14.528522

**Authors:** Javier Ruiz-Navarro, Sofía Blázquez-Cucharero, Víctor Calvo, Manuel Izquierdo

## Abstract

T cell receptor (TCR) stimulation of T lymphocytes by antigen bound to the major histocompatibility complex (MHC) of an antigen-presenting cell (APC), together with the interaction of accessory molecules, induces the formation of the immunological synapse (IS), the convergence of secretion vesicles towards the centrosome, and the polarization of the centrosome to the IS. Upon IS formation, an initial increase in cortical filamentous actin (F-actin) at the IS takes place, followed by a decrease in F-actin density at the central region of the IS, which contains the secretory domain. These reversible, cortical actin cytoskeleton reorganization processes that characterize a mature IS occur during lytic granule secretion in cytotoxic T lymphocytes (CTL) and cytokine-containing vesicle secretion in T-helper (Th) lymphocytes. Besides, IS formation constitutes the basis of a signalling platform that integrates signals and coordinates molecular interactions that are necessary for an appropriate antigen-specific immune response. In this chapter we deal with the three-dimensional (3D) analysis of the synaptic interface architecture, as well as the analysis of the localization of different markers at the IS.

## 1 Introduction

### 1.1 The immunological synapse

The IS is a specialized cell-cell interface contact area, established between a T lymphocyte and an APC carrying cognate MHC-bound antigen, that provides a signalling platform for integration of signals leading to execution of an immune response. This allows intercellular information exchange, in order to ensure efficient TCR signal transduction, T cell activation and the proper execution of diverse T lymphocyte effector functions (Dustin, 2016). IS establishment is a very dynamic, plastic and critical event, enabling the integration of spatial, mechanical and biochemical signals, involved in specific, cellular and humoral immune responses (Fooksman et al., 2010) (de la Roche et al., 2016). The general architecture of the IS upon cortical actin reorganization is recognized by the formation of a concentric, bullseye spatial pattern, named the supramolecular activation complex (SMAC) (Griffiths et al., 2010) (Billadeau et al., 2007) (Kuokkanen et al., 2015) (Yuseff et al., 2013) (Carisey et al., 2018). This reorganization yields three differentiated regions: first, a central cluster of antigen receptors bound to antigen called central SMAC (cSMAC) produced by centripetal traffic; second, a surrounding ring rich in adhesion molecules, such as the LFA-1 integrin, called peripheral SMAC (pSMAC), which appears to be crucial for adhesion with the APC (Monks et al., 1998) (Fooksman et al., 2010); third, at the edge of the contact area with the APC, the pSMAC-surrounding distal SMAC (dSMAC), which comprises a circular array of dense F-actin (Griffiths et al., 2010) (Le Floc’h and Huse, 2015) (Ritter et al., 2013) (Rak et al., 2011). More recently, several super-resolution imaging techniques have revealed that at least four discrete F-actin networks form and maintain the shape and function of this canonical IS upon TCR-antigen interaction (Hammer et al., 2018) (Blumenthal and Burkhardt, 2020).

### 1.2 Polarization of centrosome and secretory granules

IS formation induces the convergence of cellular secretion vesicles towards the centrosome and, simultaneously, the polarization of this centrosome (the major microtubule-organizing center (MTOC) in lymphocytes) towards the IS (de la Roche et al., 2016) (Huse, 2012). These traffic events, acting together, lead to polarized secretion of: extracellular vesicles and exosomes coming from multivesicular bodies (MVB) by T, B lymphocytes and natural killer (NK) cells (Peters et al., 1991) (Alonso et al., 2011) (Herranz et al., 2019) (Calvo and Izquierdo, 2020); lytic granules by CTL (Stinchcombe et al., 2006) (de la Roche et al., 2016); stimulatory cytokines by Th cells (Huse et al., 2008); lytic proteases by B lymphocytes (Yuseff et al., 2013).

### 1.3 Synaptic actin cytoskeleton regulation of secretory traffic

#### 1.3.1 Cortical actin cytoskeleton

Actin cytoskeleton plays a pivotal role from the first step of IS formation, coordinating its assembly and driving, together with microtubules, the accumulation of synaptic components that are required for T cell activation (Ritter et al., 2013). However, the involvement of F-actin in IS extends beyond the rearrangement and signalling events that take place at the first steps, since F-actin also plays a crucial role in IS maintenance and in antigen receptor-derived signaling in T and B lymphocytes, ensuring a functional immune response (Billadeau et al., 2007). Refer to the excellent reviews on the IS formed by B lymphocytes (Yuseff et al., 2009) (Yuseff et al., 2013) and T lymphocytes (Billadeau et al., 2007) (Ritter et al., 2013).

At the early phases of IS formation, a protrusive actin polymerization activity drives the radially symmetric spreading of the T cell over the surface of the APC through the generation of filopodia and lamellipodia (Le Floc’h and Huse, 2015) (Le Floc’h and Huse, 2015)6). As the IS evolves, cortical F-actin density progressively decreases in the cSMAC, while F-actin reorganizes and accumulates into the dSMAC, creating a peripheral F-actin-rich ring, and allowing the directional secretion toward the APC by congregating secretion vesicles on the IS (Stinchcombe et al., 2006). Thus, F-actin forms a permissive network at the IS of CTL and NK cells (Ritter et al., 2015) (Carisey et al., 2018).

The concentric F-actin architecture and the cortical actin cytoskeleton reorganization are shared by IS formed by B lymphocytes, CD4+ Th lymphocytes, CD8+ CTL, and NK cells (Le Floc’h and Huse, 2015) (Brown et al., 2011). Remarkably, all these immune cells form IS and directionally secrete proteases, cytokines or cytotoxic factors at the IS. This directional secretion, regulated by F-actin synaptic architecture, most probably enhances the specificity and the efficacy of the subsequent responses to these molecules (Le Floc’h and Huse, 2015), by spatially and temporally focusing the secretion at the synaptic cleft (Billadeau et al., 2007), which avoids the stimulation or death of bystander cells. However, in this paper we deal only with IS made by T lymphocytes, although the method can be extended to the IS made by the other immune cells.

#### 1.3.2 Cortical and non-cortical actin cytoskeleton regulation of secretory traffic

F-actin reduction at the cSMAC does not just allow secretion, since it apparently also plays a key role in MTOC movement towards the IS (Stinchcombe et al., 2006) (Ritter et al., 2015). IS-induced actin cytoskeleton reorganization and secretory vesicles polarized traffic are regulated by two major pathways: one involves HS1/WASp/Arp2/3 complexes acting on cortical F-actin, and the other involves formins, a conserved family of proteins, with members such as formin-like 1 (FMNL1) and Diaphanous 1 (Dia1), that nucleate and elongate unbranched actin filaments (Kumari et al., 2014; Kühn et al., 2014; DeWard et al., 2010). In this context, several studies suggest that cortical F-actin reorganization at the IS is necessary and sufficient for centrosome and secretory granule polarization (Ritter et al., 2015) (Chemin et al., 2012) (Sanchez et al., 2019). However, other results show that FMNL1 or Dia1 depletion impedes centrosome polarization without affecting Arp2/3-dependent cortical F-actin reorganization (Gomez et al., 2007) supporting that, at least in the absence of FMNL1 or Dia1, cortical F-actin reorganization is not sufficient for centrosome polarization. Conversely, the centrosome can polarize normally to the IS in the absence of cortical actin reorganization at the IS occurring in Jurkat T lymphocytes lacking Arp2/3 (Gomez et al., 2007) (Kumari et al., 2014). Thus, analysis of all the F-actin networks at different subcellular locations is necessary to achieve the full picture of the cellular actin cytoskeleton reorganization processes leading to polarized secretion. These results, together with our results showing that PKC*δ* interference affects cortical F-actin at the IS (Herranz et al., 2019), and PKC*δ* regulates the phosphorylation of FMNL1 (Bello-Gamboa et al., 2020), prompted us to study the involvement and synaptic distribution of cortical F-actin and FMNL1 formin in T lymphocyte IS structure. The experimental approach described here allows the simultaneous assessment of the synaptic architecture and colocalization of two different markers in the three dimensions of the IS. This method is properly developed for confocal, but also for super-resolution microscopy images. As already mentioned, the crucial importance of the synaptic structure in processes such as centrosome polarization or secretory vesicles traffic and degranulation has stimulated the development of accurate analysis methods of the synaptic interface. Moreover, considering that actual biological interactions occur in three-dimensional space, following this approach, we can study the architecture of the IS and the three-dimensional colocalization and potential interactions between different markers in this region, thus overcoming the limitations of two-dimensional analyses.

## 2. Materials, cells, immunological synapse formation and image capture

1. Raji B cell line is obtained from the ATCC and Jurkat T clone C3 has been described (Herranz et al., 2019).

2. Cell lines are cultured in RPMI 1640 medium supplemented with L-glutamine (Invitrogen), 10% heat-inactivated fetal calf serum (FCS) (Gibco) and penicillin/streptomycin (Gibco) and 10 mM HEPES (Lonza).

3. For IS formation, Raji cells are attached to *μ*-Slide 8 Well Glass Bottom (Ibidi), using 20 *μ*g/ml poly-L-lysine (SIGMA) at 37 °C for 1 h, and then labelled with 10 *μ*M CellTracker(tm) Blue (7-amino-4-chloromethylcoumarin, CMAC, ThermoFisher), in supplemented RPMI 1640 medium at 37 °C for 45 min, and pulsed with 1 µg/ml Staphylococcal enterotoxin E (SEE, Toxin Technology, Inc.) at 37 °C for 45 min. After removing the culture medium containing SEE, Jurkat cells are directly added to the wells, and incubated at 37 °C for 1 h, so that IS are formed. CMAC labelling allows discriminating Jurkat-Raji conjugates (Figs. 1 and 2). Please refer to the following references for further details, since this IS model has been exhaustively described (Montoya et al., 2002) (Alonso et al., 2011) (Bello-Gamboa et al., 2019). Endpoint fixation is performed with 4% paraformaldehyde at room temperature (RT) for 15 min, and subsequently with chilled acetone for 5 min (see *Note 1*).

**Figure 1.**
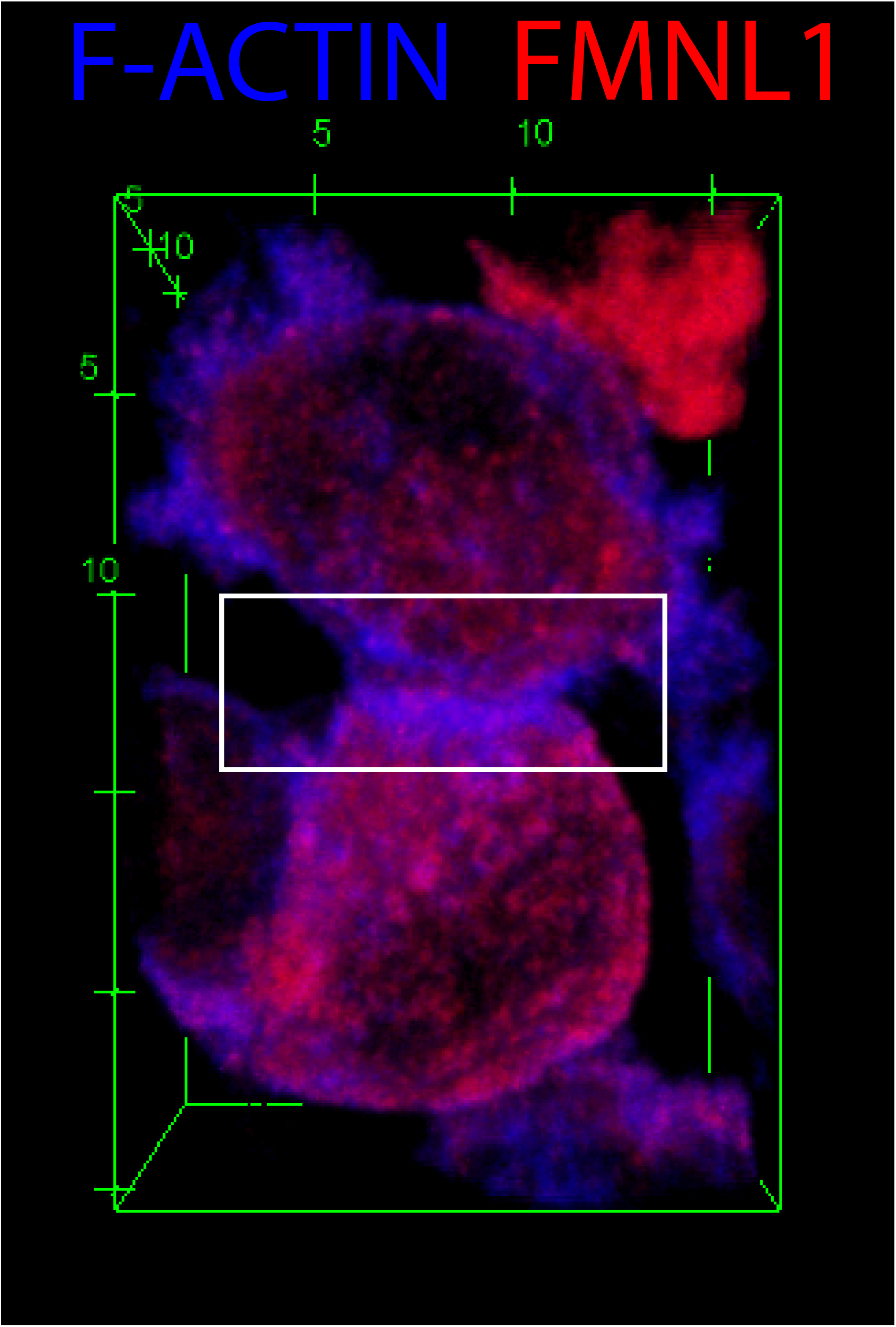
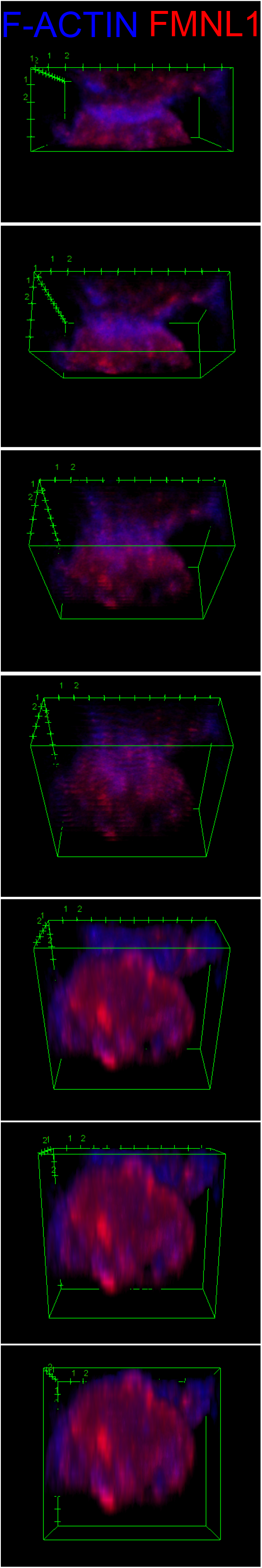
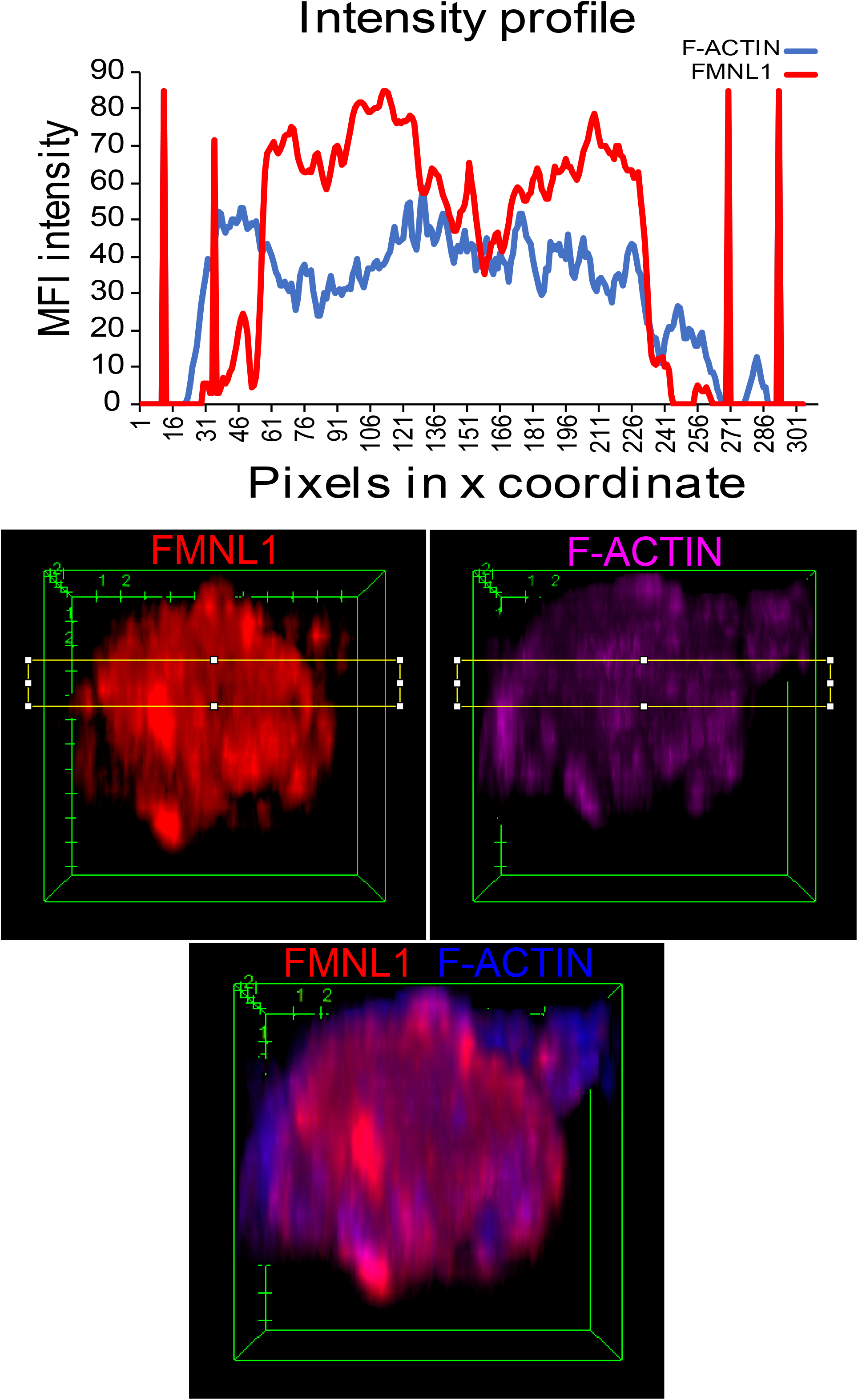
ImageJ-based 3D analyses of the immunological synapse. Jurkat cells were challenged with CMAC-labelled (blue), SEE-pulsed Raji cells for 1 h, then they were fixed, stained with anti-FMNL1 AF546 to label FMNL1 (red) and phalloidin AF647 to label F-actin (magenta) and imaged by confocal fluorescence microscopy. Panel A, top view of a whole synaptic conjugate labeled with anti-FMNL1 (red) and F-actin (blue). The white rectangle labels the crop that contains the IS area analysed in the panel B. Panel B, sequential frames (one each 20 frames) taken from Suppl. Video 1. Panel C, last frames from Suppl. Video 1 (IS interface) and corresponding mean fluorescence intensity (MFI) intensity plots of FMNL1 (red) and F-actin (blue) channels. F-actin profile reveals a multifocal IS, with several low F-actin MFI areas. In general, the F-actin density (blue line) at the central IS, that contains the cSMAC, is lower than at the edges (corresponding to the dSMAC). The FMNL1 profile (red line) is partially similar to F-actin profile.

**Figure 2.**
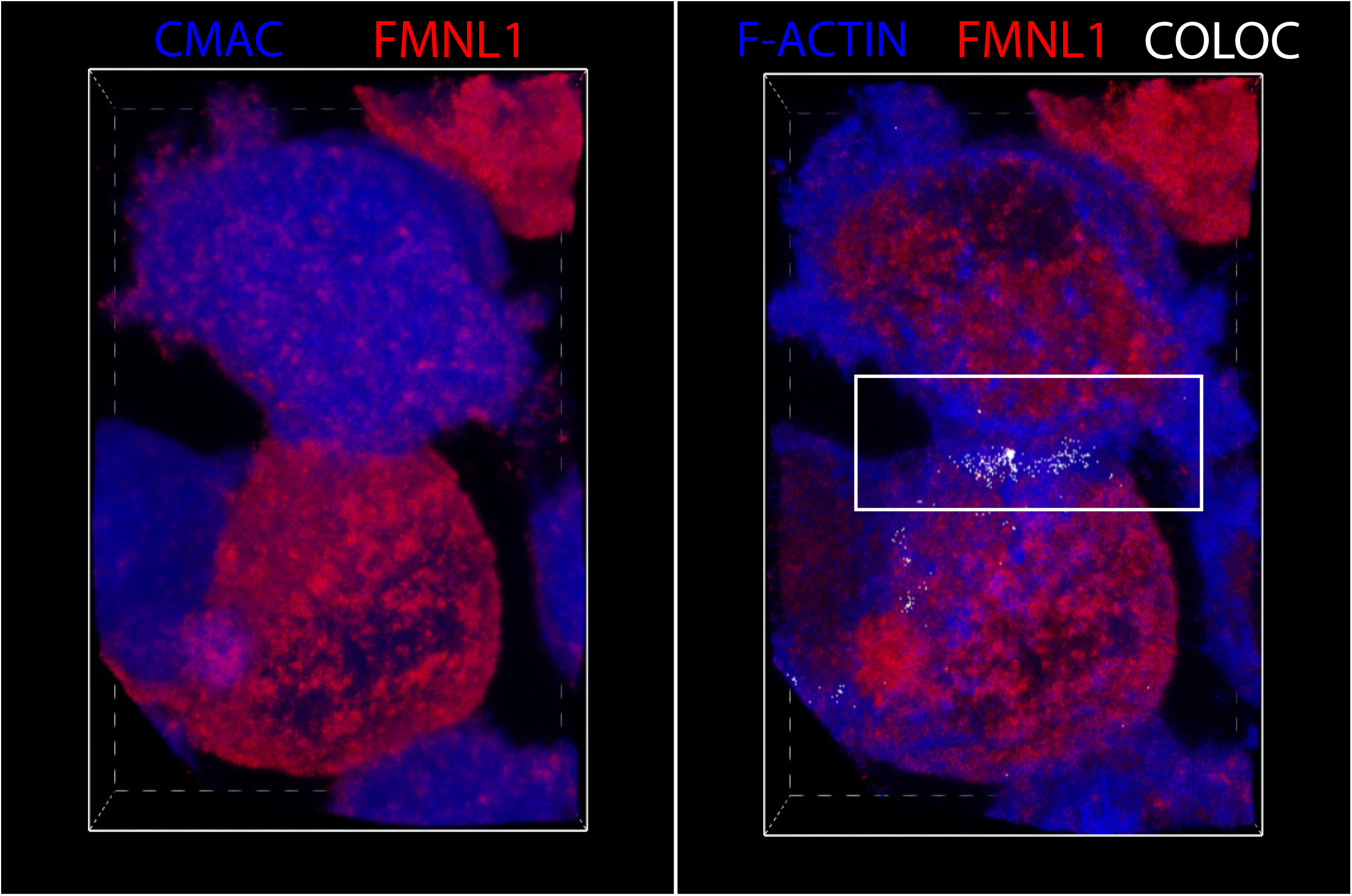
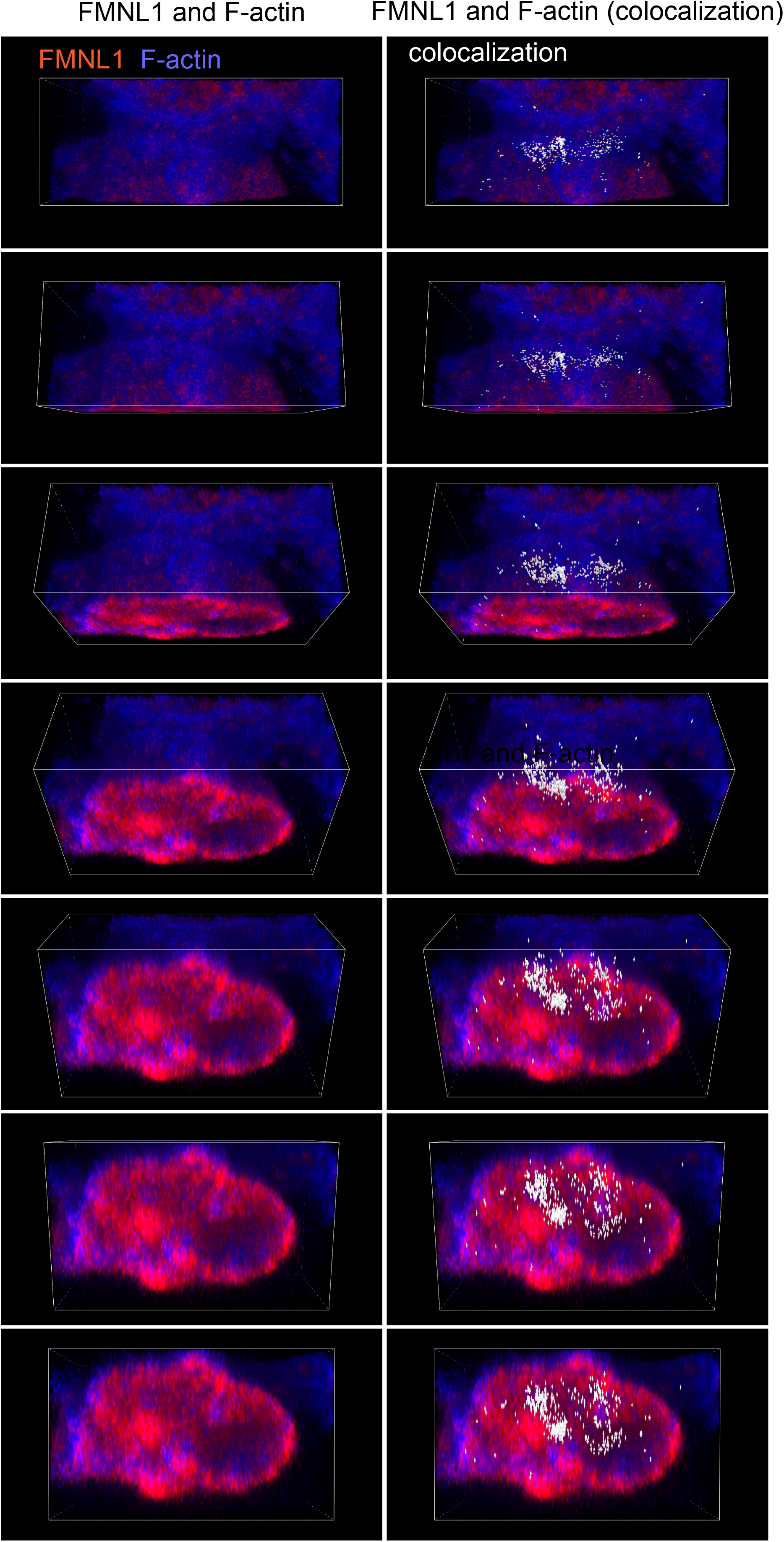
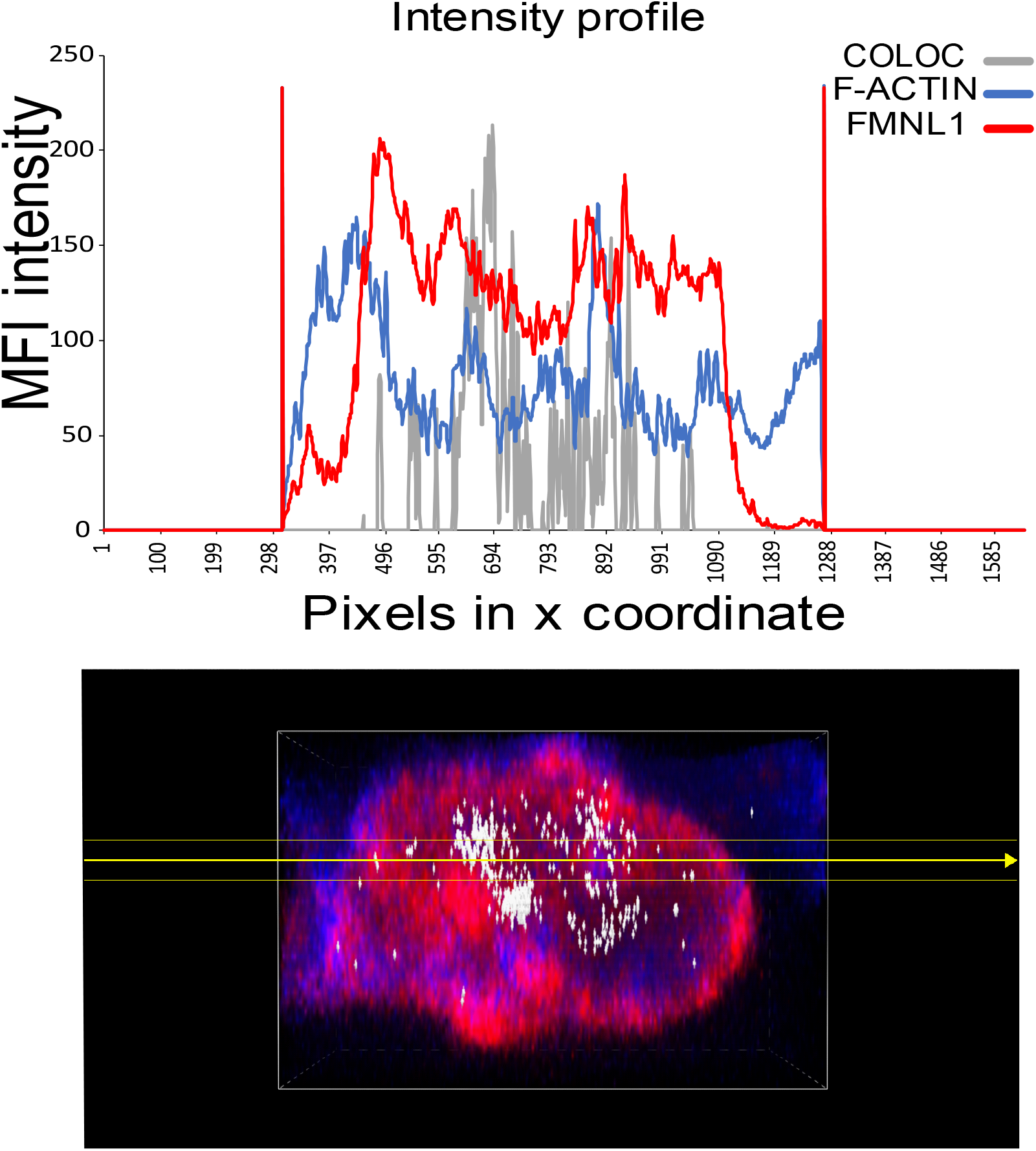
NIS-AR-based 3D analyses of the immunological synapse and 3D colocalization. Jurkat cells were challenged with CMAC-labelled (blue), SEE-pulsed Raji cells for 1 h, then they were fixed, stained with anti-FMNL1 AF546 to label FMNL1 (red) and phalloidin AF647 to label F-actin (magenta) and imaged by confocal fluorescence microscopy. In panel A, left panel shows the top view of a whole synaptic conjugate (Raji cell labeled with CMAC, blue; anti-FMNL1, red), whereas right panel displays the top view of the same synaptic conjugate showing 3D colocalization pixels between FMNL1 (red) and F-actin (blue) in white color. The white rectangle labels the crop area that contains the IS area analysed in panel B. Panel B, sequential frames (one each 20 frames) taken from Suppl. Video 2, showing 3D colocalization pixels (white color) on each frame. Colocalization pixels between FMNL1 and F-actin accumulate at the central area of the IS. Panel C, last frame from Suppl. Video 2 (IS interface) and MFI intensity plots of FMNL1 (red), F-actin (blue) channels and colocalization (grey). F-actin profile reveals a multifocal IS, with several F-actin low areas. In general, F-actin density (blue line) at the central IS, that contains the cSMAC, is lower than at the edges (corresponding to the dSMAC). FMNL1 profile is partially similar to F-actin profile, decreasing at the center of the IS, whereas the F-actin and FMNL1 colocalization (grey line) is higher at the central IS.

4. Immunofluorescence is performed as described (Fernández-Hermira et al., 2023). FMNL1 is labelled with mouse monoclonal anti-FMNL1 (clone C5, Santa Cruz Biotechnology) at RT for 45 min, and an appropriate secondary antibody coupled to AF546 (ThermoFisher) at RT for 30 min. F-actin is labelled with phalloidin AF647 (ThermoFisher) at RT for 30 min. CMAC fluorescence does not overlap with AF546 or AF647 fluorescences.

5. Image capture is performed with Leica SP8 confocal microscope. Around 30 optical sections are acquired using a 0.2-0.3 µm Z-step size to yield an appropriate axial resolution. Confocal settings are same as those previously used (Fernández-Hermira et al., 2023).

## 3. 3D analysis of the IS

### 3.1 Image setup and formatting

1. Open the confocal file in the public, multiplatform software ImageJ or FIJI (https://imagej.nih.gov/ij/, https://fiji.sc), by selecting “Open” > “Select File” > “OK”. All channels saved for each selected image should open automatically using “Bioformats import” plugin. Depending on the confocal file source it could be necessary to specify pixel width, height and voxel depth in microns. Please include in “Image” > “Properties” these parameters obtained from confocal image metadata. This also allows proper image calibration.

2. Save all individual channels in TIFF format by using “File” > “Save As” > “OK”.

### 3.2 ImageJ 3D synapse analysis

#### 1. Channel rotation

1.1. Open a TIFF file in ImageJ or FIJI by selecting “Open” > “Select File” > “OK”. All channels saved for each selected image should open automatically.

1.2. Rotate channel by using “Image” > “Transform” > “Rotate”. Enter desired angle so as to position the Raji B cell vertically to the Jurkat T cell. The synapse should appear horizontally to the observer. “Preview” function allows to view the image with the selected angle. By clicking “OK”, modifications are saved. Repeat process with the same angle for all channels.

1.3. Save all individual channels by using “File” > “Save As” > “Tiff” > “OK”.

#### 2. Channel crop

2.1. To facilitate cropping, synchronize all channels by using “Analyze” > “Tools” > “Synchronize Windows”. Select “Synchronize All” in pop-up window. Channels stay synchronized between each other as long as the pop-up window stays open.

2.2. Select area to crop using the “ rectangle” tool. Same area and same position are reproduced in all synchronized crops. Use “Image” > “Crop”. Select an area comprising both the entire Raji B and Jurkat T cells (Fig. 1A). A second crop can be generated in order to select only the synapse area between the two cells (see *Note 2*) (Fig. 1B). If channels are not synchronized, setting up the same width “w” and height “h” values, and “x” and “y” positions, ensures the equivalence in size and position between all cropped channels. Values for “w”, “h”, “x” and “y” are displayed in FIJI’s console.

2.3. Save rotated cropped channels as TIFF files.

#### 3. Generation of the 3D file and video recording

3.1. Image look up tables (LUTs) can be adjusted by using “Image” > “Adjust” > “Brightness/Contrast”.

3.2. Merge two or more channels by using “Image” > “Color” > “Merge channels”. Choose color components for each channel and select “OK” (see *Note 3*). Select “Keep source images” to keep original images.

3.3. Create a 3D image by selecting “Plugins” > “3D Viewer”. In “Image”, select desired file for video. Leave other parameters as given. Click “OK”.

3.4. To visualize the delimitating box for the video, select “Edit” > “Show bounding box”.

3.5. To generate video, select “View” > “Start freehand recording” and create sequential frames by using the arrows in the keyboard (see *Note 4*). Alternatively, select “Set View” to choose the desired plane.

3.6. To stop recording, select “View” > “Stop freehand recording”. A window named “Movie” appears.

3.7. “Movie” file can be saved by using “File” > “Save As” > “AVI” (Fig. 1). Once the .avi stack is generated (Supplementary Video 1) the video stack can be converted to sequential images by using “Image” > “Stacks” > “Stack to images” (Supplementary Video 1, Fig. 1B).

#### 4. Quantification and measurement of IS interface (en face) intensity profile

4.1. In order to obtain a synapse’s intensity profile for each channel, individual recordings should be generated for all channels included in the video.

4.2. Scroll forementioned videos to those frames where the interface of the synapsis is shown perpendicular (en face) to the observer.

4.3. Select area over the en face frame to create the intensity profile. Save this by using “Analyze” > “Tools” > “ROI Manager”. Click on “Add” and “More” > “Save”.

4.4. To obtain the intensity profile graph, select “Analyze” > “Plot Profile”. Save each individual channel graph by using “File” > “Save As” > “Tiff”.

4.5. To obtain numerical data, select “List” in graph and copy the data onto an Excel spreadsheet. By copying the individual data from all channels and inserting the desired graph, a new intensity profile plot can be generated that emulates all chosen channels (Fig. 1C).

### 3.3 NIS-Elements AR 3D synapse analysis

#### 1. Channel rotation

1.1. Open the software NIS-Elements AR (https://www.microscope.healthcare.nikon.com/en_EU/products/software/niselements/nis-elements-advanced-research) (NIS-AR) and a TIFF confocal file generated in step 1.1 by using “Image” > “Open” > “OK”.

1.2. Rotate channel by using “Image” > “Rotate” > “Rotate by Degree”. Enter desired angle so as to position the Raji B cell vertically to the Jurkat T cell. The synapse should appear horizontally to the observer.

1.3. Repeat process with the same angle rotation for all channels. Save rotated channels as separated, individual TIFF files by selecting “File” > “Save/Export to TIFF Files” > “Choose Output Folder” > “Export”.

#### 2. Channel crop

2.1. Open rotated TIFF channels and select “Image” > “Crop”. Select area comprising both the entire Raji B and Jurkat T cells (Fig. 2A). A second crop can be generated in order to show only the synapse between the two cells (see *Note 2*) (Fig. 2A and B). By ticking “Remember Last Settings” tool, all subsequent crops should measure the same and be located at the same x, y position.

2.2. Repeat process for all rotated channels. Save rotated cropped channels as separated, individual TIFF files by selecting “File” > “Save/Export to TIFF Files” > “Choose Output Folder” > “Export”.

#### 3. Generation of the 3D file

3.1. Open two equally edited channel TIFF files by using “File” > “Open” > “OK”.

3.2. The “Experiment Type” window appears automatically. Enter Z series width for each channel.

3.3. Merge both channels by using “File” > “Merge Channels”. Choose RGB color components for each channel (see *Note 3*).

3.4. Image LUTs can be adjusted by using “Visualization Controls” > “LUTs”.

3.5. Colocalization analysis.

- Show fluorocytogram by selecting “View” > “Analysis Controls” > “Colocalization”. Define colocalization region with “Rectangular Selection” or “Sector Selection” symbols (see *Note 5*).
- Change colocalization color to white, “View” > “Analysis Controls” > “Binary Layers” > “Colocalization” > “More” > “Select desired color” > “Accept”, to distinguish better the colocalization pixels from the other channel colors.
- Save the colocalization selected region by selecting “Save/Load Colocalize Configuration” symbol > “Save Colocalization Settings” > “Save”.

3.6. Create a 3D image by selecting “Show Volume View” symbol on the top toolbar (Fig. 2B).

#### 4. Video recording of the 3D file

4.1. While on the 3D generated image, open the video editor with “Show Movie Maker” tool on the top toolbar.

4.2. Select the desired starting frame by using manual tools or by selecting “View Plane” on the top toolbar and choosing the desired plane. Select “Plus” symbol on the bottom of the screen to fix initial settings.

4.3. Repeat action as many times as needed to create the final movie. Added blocks can be individually edited by right-clicking on them.

4.4. Select “Play/Stop Movie” to show the recording.

4.5. Save selected recording by using “Export Movie” symbol on the bottom of the screen. Save the generated recording by clicking “File” > “Save As” both in .nd2 and .avi formats for future editing and viewing. Once the .avi stack is generated (Supplementary Video 2) the video stack can be converted to serial images by using “Image” > “Stacks” > “Stack to images” (Fig. 2B).

#### 5. Quantification and measurement of synapse’s en face intensity profile

5.1. Display the intensity profile tool in NIS-AR.

5.1. Open the desired video in .nd2 format by using “File” > “Open”.

5.3. Scroll the video to the time frame where the IS interface is shown perpendicular to the observer (en face) by reproducing the video through all its time frames using the bottom blue T level bar.

5.4. Select the “Intensity Profile” symbol on the top toolbar and adjust the Profile Line by dragging it until it is positioned in the desired area (to analyze colocalization data, see *Notes 6* and *7*). The x, y coordinates of the Profile Line can be viewed at the bottom of the screen, and used, if necessary, in other videos.

5.5. Upon generation of the intensity profile graph, separate profiles of each selected channels and the colocalization channel are shown in the “Intensity Profile” window

5.6. Export the data to an Excel spreadsheet by selecting “Export” on the “Intensity Profile” window. A new Excel document is automatically generated with the raw data. By selecting the appropriate data and inserting the desired graph, a new separate Intensity Profile graph for each channel, including the colocalization channel, can be generated (Fig. 2C).

## 4. Discussion, future perspectives and concluding remarks

We have noticed that the quality of the 3D videos generated by NIS-AR is higher than that of videos generated using ImageJ/3D viewer, and that both lagging and poor resolution can be appreciated in some frames of ImageJ-generated videos. In addition, NIS-AR software easily handles the files, whereas we have observed certain software instability during ImageJ processing in three different computers. In addition, 3D colocalization plugin is not available in ImageJ yet, please check for plugin actualizations in ImageJ website (https://imagej.nih.gov/ij/). It is important to remark that the procedure described here is intended to handle post-processing of confocal imaging in fixed samples, but there are some alternatives for 3D reconstitution and IS interface visualization of the IS in living cells with high temporal resolution, albeit the equipment involved is scarce and expensive (Calvo and Izquierdo, 2018). These techniques allow image analyses of developing synapses carried out during the acquisition stage, by using light sheet fluorescence microscopy (LSFM) (Ritter et al., 2015) (Calvo and Izquierdo, 2018). Indeed, LSFM technique allows four-dimensional (x, y, z, t) analyses of IS interface and F-actin architecture using Sir Actin probes in living cells, but the superresolution option is not available at this moment using this technique. In addition, other approaches based on stimulatory artificial surfaces (lipid bilayers) and total internal reflection microscopy (TIRFM), combined with superresolution techniques such as structured illumination microscopy (SIM) allow superresolution images of IS interface and superb F-actin imaging in living synapses, by reducing 3D dimensions to two (Calvo and Izquierdo, 2018) (Murugesan et al., 2016) (Carisey et al., 2018), although these approaches are intrinsically reductionist and do not mimic the complex interactions and the irregular stimulatory surface taking place in a “real” cell-cell synapse (Dustin, 2009) as we present here.

As described, the protocols can be used to handle files generated in diverse confocal microscopes due to the flexibility of ImageJ. Thus, it is conceivable that the resolution and quality of the generated videos will greatly improve with the enhanced resolution and quality of the source images, including superresolution images (Calvo and Izquierdo, 2018).

The method described above is not automated and to perform the quantification the user must therefore select single cells forming IS one by one, which may make it difficult to evaluate results in a fully unbiased manner and may also be time-consuming to execute for an untrained user. However, a trained user may analyze up to 4-5 synapses per hour. To prevent biases, please try to randomly select as many synaptic conjugates from different fields as possible. Unfortunately, it is not easy to distinguish canonic IS from irrelevant cell contacts or cell aggregates randomly produced by the high cell concentrations used to favor IS formation (Bello-Gamboa et al., 2019). Evaluation of the cup-shaped profile of the effector T lymphocyte and/or lamellipodium formation observed in the transmittance channel may help to identify IS (Fernández-Hermira et al., 2023). In addition, a useful criterion to unambiguously select canonic IS and differentiate them from irrelevant cell contacts is to use the F-actin probe phalloidin (Fig. 1 and Fig. 2), since it is known that F-actin accumulates at the lamellipodium in the synaptic contact area (see 1. Introduction). CMAC labelling facilitates discrimination of Jurkat-Raji conjugates from Jurkat-Jurkat or Raji-Raji homotypic conjugates.

In addition, and due to the fact that the IS characteristic concentric bullseye F-actin architecture and the actin cortical cytoskeleton reorganization processes are shared by IS formed by B lymphocytes, CD4+ Th lymphocytes, CD8+ CTL, and NK cells forming IS (Le Floc’h and Huse, 2015) (Brown et al., 2011), the protocol described here can be extended to analyze IS formed by all these types of immune cells. This analysis will enable to address how the different cortical F-actin network areas in all these IS classes are regulated, how these networks integrate into cell surface receptor-evoked signalling networks, as well as their potential regulators, that constitute an intriguing and challenging biological issue (Dogterom and Koenderink, 2019) (Calvo and Izquierdo, 2021).

## 5. Notes

1. Acetone (and acetone vapor) dissolves plastic, thus keep the *μ*-Slide with fixative on ice at 4 °C, and remove the *μ*-Slide plastic lid. Recommended maximal fixation time is 5 min, since longer times dehydrate samples in excess (producing cell shrinking and cell shape changes). Paraformaldehyde or glutaraldehyde fixatives alone do not allow a clear staining of FMNL1. An important issue is that F-actin staining with phalloidin is not compatible with methanol fixing, whereas it is compatible with paraformaldehyde and/or acetone fixation. Thus, we have chosen paraformaldehyde/acetone fixation since it constitutes a good compromise to circumvent these caveats (Abrahamsen et al., 2018).

2. It is important to check all Z optical sections so as to avoid cropping out an area that includes part of one or both cells. Cropped areas can be adjusted to include the entirety of the cell area at different Z optical sections. The crops of all channels of each image must be equal both in size and x,y position.

3. For F-actin and FMNL1 staining experiment, Blue and Red Components were chosen, respectively.

4. The speed at which the keyboard arrows are clicked does not affect the final speed at which the video is recorded.

5. Once the colocalization settings are defined, load them for all Zs by scrolling the image stack through all its Z optical sections, using either the scroll wheel on the mouse or dragging them from the bottom blue Z level bar. If not, it is possible that in the subsequent 3D visualization the updated colocalization data will not be loaded.

6. Due to the properties of the video editor and the three-color nature of the “Merge Channels” tool, the generated videos will by default be RGB. For that reason, caution should be taken on the intensity profile measurements made with colocalization videos as, depending on the selected colocalization color, most probably some color components will not be properly separated in the intensity profile. For example, if, as suggested above, colocalization is displayed in white, every signal from our markers will be contaminated by the colocalization measurement. In order to solve this, a second video with the same image, but without the colocalization channel, should be created. The two different videos can be plotted in the same intensity profile graph by selecting the specific color components: for the video with colocalization the color component will be the one associated with colocalization, whereas for the video without the colocalization the color components will be all of them.

7. On the Intensity Profile graph, the signal region is flanked by two intense and very well-defined lines (Fig. 2C). These lines correspond to the boundaries of the “box” containing either the whole cell conjugate or the synapse area in the 3D display. To properly compare the Intensity Profile of different 3D videos of the same cell – as in the colocalization case – the profile line should be positioned at the same x, y coordinates in every video and the width of the generated profiles should be the same. If the 3D videos display different widths, a careful look into the sizes of the two created videos should be taken.

## Abbreviations

APC: antigen-presenting cell;
BCR: B-cell receptor for antigen;
CMAC: CellTracker(tm) Blue (7-amino-4-chloromethylcoumarin);
cSMAC: central supramolecular activation cluster;
CTL: cytotoxic T lymphocytes;
Dia1: diaphanous-1;
dSMAC: distal supramolecular activation cluster;
F-actin: filamentous actin;
FCS: fetal calf serum;
FMNL1: formin-like 1;
IS: immunological synapse;
LSFM: light sheet fluorescence microscopy;
LUTs: look up tables;
MFI: mean fluorescence intensity;
MHC: major histocompatibility complex;
MTOC: microtubule-organizing center;
MVB: multivesicular bodies;
NIS-AR: NIS-Elements
AR; NK: natural killer;
PKC*δ*: protein kinase C *δ* isoform;
pSMAC: peripheral supramolecular activation cluster;
ROI: region of interest;
RT: room temperature;
SEE: Staphylococcal enterotoxin
E; SIM: structured illumination microscopy;
SMAC: supramolecular activation cluster;
TCR: T-cell receptor for antigen;
Th: T-helper;
TIRFM: total internal reflection microscopy;
TRANS: transmittance;
3D: three dimensional.

## Author Contributions

J.R. and S.B. wrote the manuscript and prepared the figures. V.C. and M.I. conceived the manuscript and the writing of the manuscript and approved its final content. Conceptualization, V.C. and M.I; writing original draft preparation, M.I.; reviewing and editing, V.C. and M.I.

## Acknowledgements

This research was funded by grants from the Spanish Ministerio de Ciencia e Innovación, (PID2020-114148RB-I00) to MI, which was in part granted with FEDER funding (EC), corresponding to Programa Estatal de Generación de Conocimiento y Fortalecimiento Científico y Tecnológico del Sistema de i+d+i y de i+d+i Orientada a los Retos de la Sociedad.

## Figure legends

**Supplementary Video 1**

IS was performed as described in the Section 2., step 3., and the Fig. 1 legend. The video (8 frames per second) was generated as described in Section 3.2, step 3, and represents, in the initial frame, the top view of synapse area of an IS conjugate labeled with anti-FMNL1 (red) and F-actin (blue), whereas the last frame corresponds to the en face view of the IS interface.

**Supplementary Video 2**

IS was performed as described above. Both videos (24 frames per second) were generated as described in Section 3.3, step 4., and represents, in the initial frame, the top view of synapse area of an IS conjugate labeled with anti-FMNL1 (red) and F-actin (blue), whereas the last frame corresponds to the en face view of the IS interface. The lower video includes, in white color, the colocalization pixels between FMNL1 and F-actin.

